# Preventive efficacy of a tenofovir alafenamide fumarate nanofluidic implant in SHIV-challenged nonhuman primates

**DOI:** 10.1101/2020.05.13.091694

**Authors:** Fernanda P. Pons-Faudoa, Antons Sizovs, Kathryn A. Shelton, Zoha Momin, Lane R. Bushman, Jiaqiong Xu, Corrine Ying Xuan Chua, Joan E. Nichols, Trevor Hawkins, James F. Rooney, Mark A. Marzinke, Jason T. Kimata, Peter L. Anderson, Pramod N. Nehete, Roberto C. Arduino, Mauro Ferrari, K. Jagannadha Sastry, Alessandro Grattoni

## Abstract

Pre-exposure prophylaxis (PrEP) using antiretroviral oral drugs is effective at preventing HIV transmission when individuals adhere to the dosing regimen. Tenofovir alafenamide (TAF) is a potent antiretroviral drug, with numerous long-acting (LA) delivery systems under development to improve PrEP adherence. However, none has undergone preventive efficacy assessment. Here we show that LA TAF using a novel subcutaneous nanofluidic implant (nTAF) confers partial protection from HIV transmission. We demonstrate that sustained subcutaneous delivery through nTAF in rhesus macaques maintained tenofovir diphosphate concentration at a median of 390.00 fmol/10^6^ peripheral blood mononuclear cells, 9 times above clinically protective levels. In a non-blinded, placebo-controlled rhesus macaque study with repeated low-dose rectal SHIV_SF162P3_ challenge, the nTAF cohort had a 62.50% reduction (95% CI: 1.72% to 85.69%; *p*=0.068) in risk of infection per exposure compared to the control. Our finding mirrors that of tenofovir disoproxil fumarate (TDF) monotherapy, where 60.00% protective efficacy was observed in macaques, and clinically, 67.00% reduction in risk with 86.00% preventive efficacy in individuals with detectable drug in the plasma. Overall, our nanofluidic technology shows potential as a subcutaneous delivery platform for long-term PrEP and provides insights for clinical implementation of LA TAF for HIV prevention.

## 1. Introduction

The approval of Descovy® (200 mg emtricitabine [FTC]/25 mg tenofovir alafenamide [TAF]) as the second HIV pre-exposure prophylaxis (PrEP) medication, following Truvada® (200 mg FTC/300 mg tenofovir disoproxil fumarate [TDF]) is fueling global efforts to end the AIDS pandemic by 2030.^[1]^ Compared to Truvada®, Descovy® offers safety advantages with lower systemic tenofovir (TFV) concentrations without compromising overall efficacy (NCT02842086).^[2]^ The efficacy of these agents to prevent sexual HIV infection is exceptional, provided that individuals strictly adhere to the dosing regimen.^[3–5]^ According to the iPrEx study, seven doses of Truvada® per week correlated with 99% PrEP efficacy, whereas the rate dropped to 76% with two doses per week.^[6]^ Motivated by challenges of pill fatigue and PrEP accessibility, various biomedical developments have emerged aiming at improving therapeutic adherence and expanding HIV PrEP implementation.

Long-acting (LA) antiretroviral (ARV) formulations and delivery systems offer systemic delivery for prolonged periods, obviating the need for frequent dosing. Currently, LA ARV strategies for HIV PrEP are largely geared towards developing single-agent drugs for prevention instead of combinatorial formulations.^[7–15]^ Focusing on a single drug allows for maximal drug loading, while minimizing injection volumes (for injectables). In the case of LA ARV implants, a single drug formulation affords smaller size dimensions for minimally-invasive and discreet implantation.^[16, 17]^ Importantly, single-agent LA ARVs offer benefits of cost-effectiveness as well as reduced complexity in terms of development. Of relevance, a single-agent injectable LA ARV, cabotegravir, is currently in clinical trials for PrEP efficacy evaluation (NCT02076178, NCT02178800, NCT02720094, NCT03164564).^[18, 19]^ Thus far, islatravir (MK-8591) remains the only single-agent ARV LA ARV implant to reach clinical testing for safety and pharmacokinetics assessment.^[20]^

Given the potency and safety advantages of TAF compared to TDF, numerous LA TAF strategies are under development involving biodegradable^[7–9]^ or non-biodegradable^[10]^ polymeric implants, transcutaneously refillable devices^[11]^, and an osmotic pump system.^[15]^ While some LA TAF systems have achieved targeted preventive tenofovir diphosphate (TFV-DP) concentrations in peripheral blood mononuclear cells (PBMC) (40.0 fmol/10^6^ cells)^[7,10,11]^, none has undergone efficacy studies for protection from HIV transmission. Thus, considering the concentrated research efforts on developing LA TAF systems, it is of utmost importance to evaluate the efficacy of LA TAF as a single-agent drug for HIV prevention.

Here, we present the first efficacy study of LA TAF for HIV PrEP. We used a nonhuman primate (NHP) model of repeated low-dose rectal challenge with simian HIV_SF162P3_ (SHIV_SF162P3_), which recapitulates human HIV transmission. We assessed the efficacy of sustained subcutaneous delivery of TAF via a novel nanofluidic (nTAF) implant as a single-agent PrEP regimen for protection from SHIV_SF162P3_ infection. We investigated the pharmacokinetics and biodistribution of TAF, as well as safety and tolerability of the implant.

## 2. Results

### 2.1. Nanofluidic implant assembly

We leveraged a newly designed silicon nanofluidic membrane technology^[21]^ for sustained drug elution independent of actuation or pumps. The nanofluidic membrane (6 mm × 6 mm with a height of 500 μm) is mounted within a medical-grade titanium drug reservoir **(Figure 1A)**. The nanofluidic membrane contains 199 circular microchannels, each measuring 200 μm in diameter and 490 μm in length. Hexagonally distributed in a circular configuration (Figure 1B), each microchannel leads to 1400 parallel slit-nanochannels (Figure 1C), for a total of 278,600 nanochannels per membrane. The nanochannels (length 10 μm, width 6 μm) are densely packed in square arrays organized in circular patterns. The whole membrane surface is coated by an innermost layer consisting of silicon dioxide (SiO_2_), and a surface layer of silicon carbide (SiC), which provides biochemical inertness for long term implantable applications (Figure 1D).^[22,23]^

**Figure 1.**
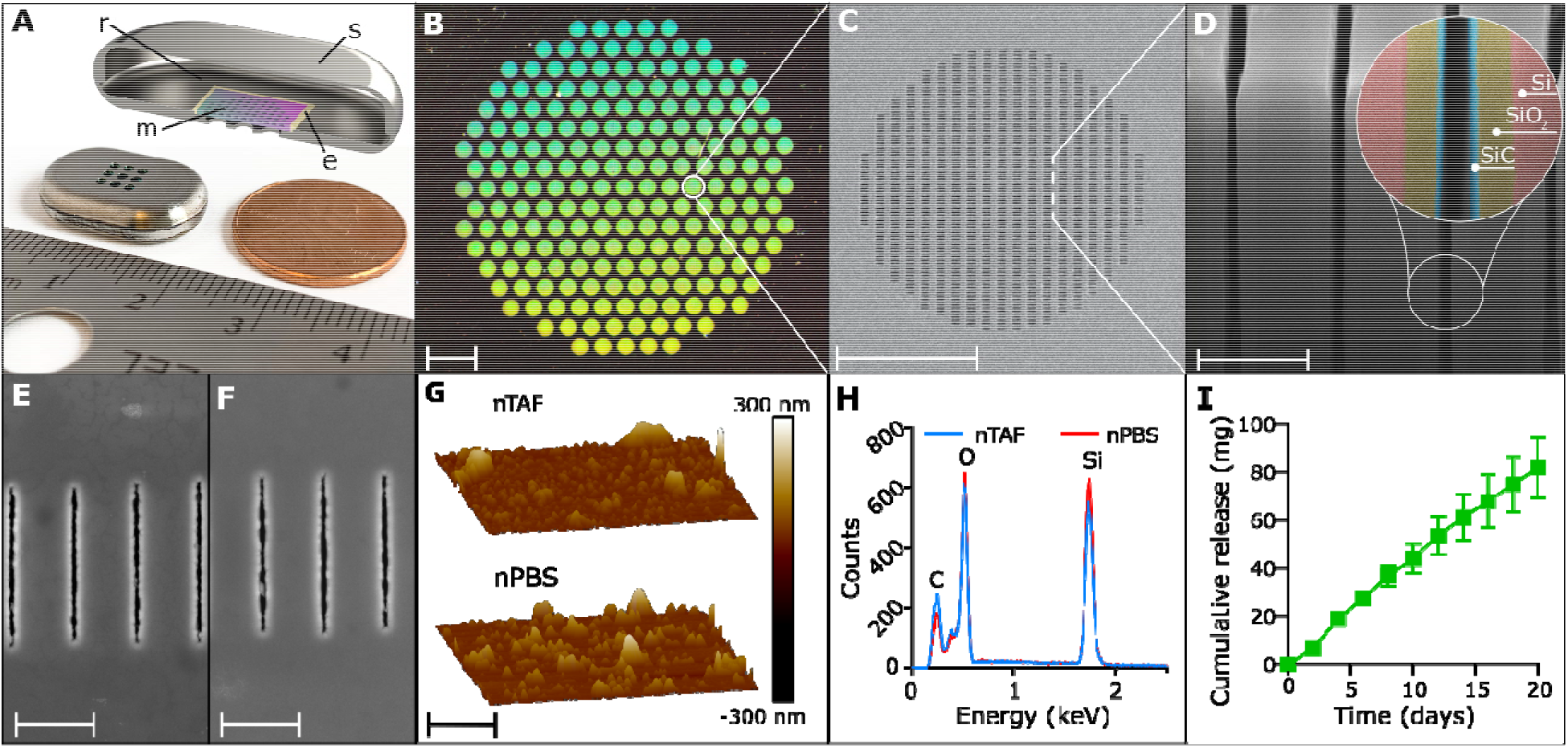
The nanofluidic implant for subcutaneous TAF HIV PrEP delivery. A) Rendered image of cross-section of titanium drug reservoir. B) Assembled titanium TAF drug reservoir with 200 nm nanofluidic membrane. Image taken at 0.5 x magnification, scale bar is 1 mm. C) Top-view of SEM image of nanochannel membrane. Scale bar is 100 μm. D) FIB image of nanochannel membrane cross-section displaying perpendicular nanochannels. Zoom-in on nanochannel layers colored for identification. Scale bar is 2 μm. E) Representative top view SEM image of nanochannel membrane from nTAF after 4 months in vivo. Scale bar is 2.5 μm. F) Representative top view SEM image of nanochannel membrane from nPBS after 4 months in vivo. Scale bar is 2.5 μm. G) Representative AFM image of membrane from nTAF compared to AFM image of membrane from nPBS after 4 months in vivo. Scale bar is 2.5 μm. H) EDX analysis of surface elements below SiC coating of membrane from nTAF compared to nPBS after 4 months in vivo. I) Cumulative release of drug *in vitro* (mean ± SEM) from nTAF into sink solution (n=5). SiC, silicon carbide, SiO_2_, silicon oxide, Si, silicon.

Drug diffusion across the membrane is driven by concentration difference between the drug reservoir and the subcutaneous space. The drug is loaded in the implant in powder form and is continuously solubilized in the interstitial fluids penetrated within the implant via capillary wetting of the membrane. Drug release is determined by both nanochannels and drug solubilization kinetics. Within the nanochannels, diffusivity of drug molecules is defined by steric and electrostatic interactions with channel walls. The size of nanochannels is selected to saturate drug transport, rendering it steady and independent from the concentration gradient.^[24, 25]^ The release rate can be finely tuned by selecting the suitable number of nanochannels per membrane.^[26]^ Therefore, the nanofluidic membrane passively achieves constant and sustained drug delivery obviating the need of mechanical components.^[27,28]^

In this study, based on the molecular size and physicochemical properties of TAF, we used the nanochannels size of ~190 nm. PrEP implants were loaded with solid powder TAF (nTAF), while control implants were loaded with phosphate buffered saline (nPBS). Membrane stability was evaluated after 4 months of subcutaneous implantation via scanning electron microscopy (SEM) (Figure 1E and F) along with atomic force microscopy (AFM) (Figure 1G) and energy dispersive x-ray spectroscopy (EDX) (Figure 1H). We observed similar surface morphology by AFM for the nTAF and nPBS membranes, with a non-statistically significant increase in roughness in the nPBS membrane. The EDX showed the same abundance of elements at the surface in both membranes, indicating that TAF does not alter the membrane composition. These results demonstrate that TAF does not affect membrane stability even after prolonged implantation.

Short-term *in vitro* drug release from nTAF showed a linear cumulative release of 81.85 ± 12.55 mg (mean ± SEM) of TAF over 20 days (Figure 1I). However, an increase of TAF degradation products was observed throughout the study, attributable to decrease in TAF stability (Figure S1, Supporting Information).

### 2.2. nTAF pharmacokinetic profile in NHP

For *in vivo* evaluation of pharmacokinetic (PK) and PrEP efficacy, rhesus macaques were subcutaneously implanted with either nTAF (n=8) or control nPBS (n=6) in the dorsum for 4 months. We used TFV-DP concentration in PBMC of 100.00 fmol/10^6^ cells as the benchmark prevention target, which exceeds the clinically protective level in the iPrEX trial.^[6, 7]^Preventive TFV-DP PBMC concentrations were surpassed one day post-implantation (median, 213.00 fmol/10^6^ cells; IQR, 140.00 to 314.00 fmol/10^6^ cells) and maintained at a median of 390.00 fmol/10^6^ cells (IQR, 216.50 to 585.50 fmol/10^6^ cells) for 4 months (Figure 2A). During the washout period, TFV-DP PBMC concentrations decreased to below the limit of quantitation (BLOQ) within 6 weeks of device retrieval.

**Figure 2.**
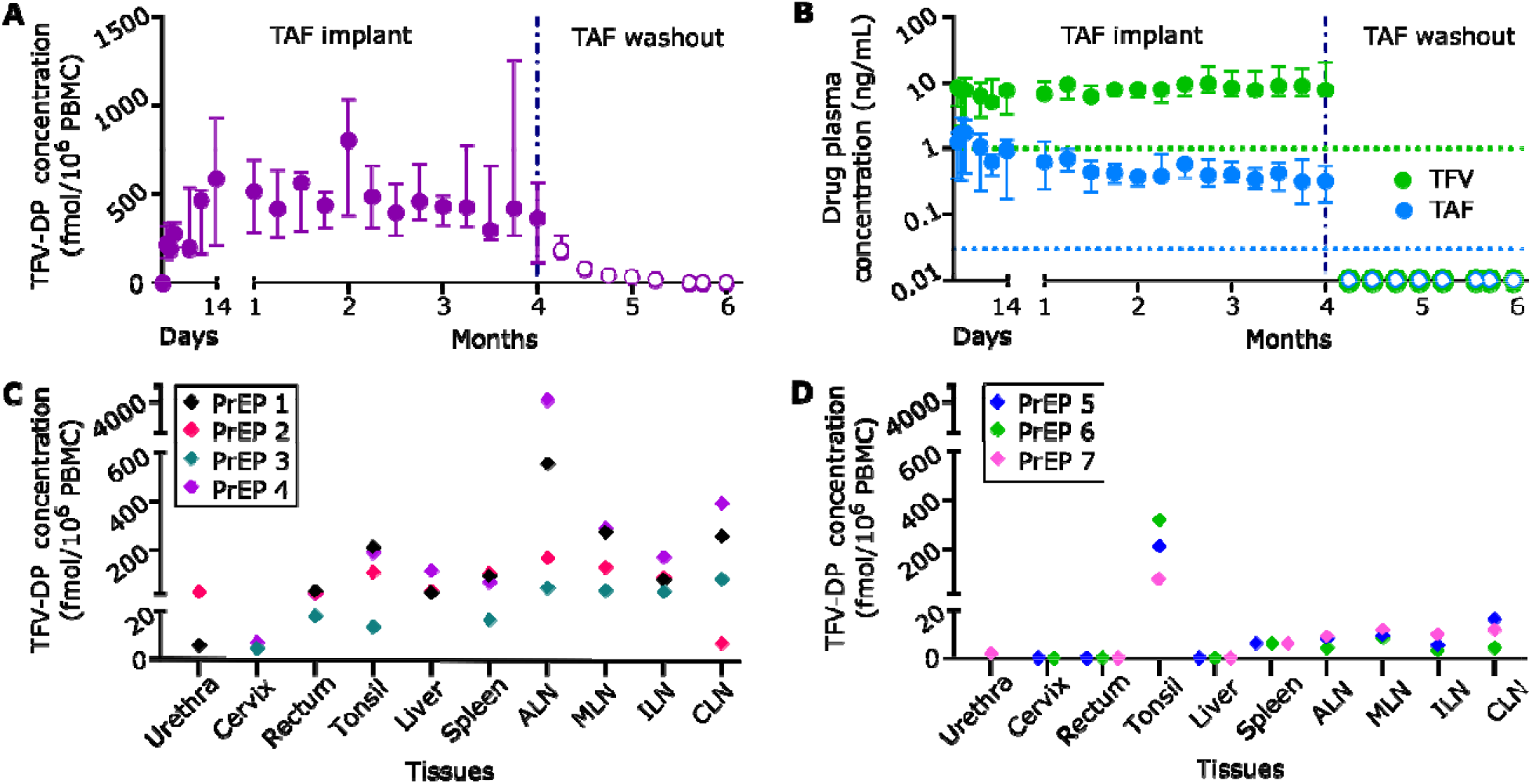
Pharmacokinetics and tissue distribution of TAF from PrEP group implanted with subcutaneous nTAF. nTAF implants (n=7) were retrieved after 4 months and washout concentrations (open circles) were followed in 3 animals. A) Intracellular TFV-DP PBMC concentrations of PrEP cohort throughout the study. B) TAF and TFV concentrations in the plasma of PrEP cohort throughout the study. Green and blue dotted horizontal lines represent lower LOQ TFV and TAF concentrations, 1.00 ng/mL and 0.03 ng/mL, respectively. C) Tissue TFV-DP concentrations upon nTAF removal after 4 months of implantation in a subset of animals (n=4). D) Tissue TFV-DP levels after the 2-month washout period in a subset of animals (n=3). Data are presented as median ± IQR in panels A and B.

Plasma TFV concentrations were consistently higher than plasma TAF for the duration of the PK study (Figure 2B). Notably, TFV concentrations increased as TAF concentrations decreased, beginning at the 3-month time point. This is attributable to the limited stability of TAF and degradation to TFV within the implant, as was observed *in vitro* (Figure S1, Supporting Information).^[29]^ Plasma TAF and TFV levels (median, 0.51; IQR, 0.30 to 0.91 ng/mL; and median, 7.81; IQR, 6.17 to 9.97 ng/mL, respectively) were within range of that achieved with oral TAF dosing of NHP.^[30]^ Within a week post-device retrieval, TAF and TFV concentrations were BLOQ.

Estimated half-life (t_1/2_) PK of TAF and TFV were below 1.87 ± 0.32 and 1.84 ± 0.63 days, respectively, as BLOQ was achieved in under a week (Table 1). Individual TFV-DP concentrations for each animal were fitted to an intravenous bolus injection two-compartment model (Figure S2A-D, Supporting Information). During the washout period, TFV-DP PBMC concentrations had an average first-order elimination rate constant of 0.14 ± 0.028 days^−1^.

**Table 1.** Plasma TAF and TFV half-lives and PBMC TFV-DP elimination rate constant pharmacokinetics in nTAF washout NHPs.

| Analyte | NHP<br>PrEP<br>5 | NHP<br>PrEP<br>6 | NHP<br>PrEP<br>7 | Average | Standard<br>deviation |
| --- | --- | --- | --- | --- | --- |
| Plasma TAF $t_{1/2}$ (days) | <2.24 | <1.71 | <1.67 | <1.87 | $\pm 0.32$ |
| Plasma TFV $t_{1/2}$ (days) | <2.55 | <1.61 | <1.35 | <1.84 | $\pm 0.63$ |
| PBMC TFV-DP $k_{10}$ (1/day) | 0.18 | 0.13 | 0.13 | 0.14 | $\pm 0.028$ |

We measured TFV-DP concentrations after device retrieval (n=4) (Figure 2C) and after the washout period (n=3) (Figure 2D) in tissues relevant to HIV-1 transmission or viral reservoirs. Specifically, we assessed cervix, urethra, rectum, tonsil, liver, spleen, axillary lymph nodes (ALN), mesenteric lymph nodes (MLN), inguinal lymph nodes (ILN), and cervical lymph nodes (CLN). Drug penetration from subcutaneous TAF delivery was observed at varying levels in all tissues after device retrieval (Figure 2C). After the two-month washout period, TFV-DP concentrations were quantifiable in the tonsil, spleen and lymph nodes (Figure 2D) and BLOQ in tissues highly associated with HIV-1 transmission, specifically the cervix and rectum. TFV-DP concentrations in the tonsil were above 75.00 fmol/mg, suggestive of longer clearance or better penetration.

### 2.3. nTAF efficacy protection against virus

We next assessed whether sustained nTAF delivery as a subcutaneously delivered monotherapy could protect the macaques against rectal SHIV_SF162P3_ infection. Prior to rectal challenge, the animals were subjected to a two-week “conditioning phase” (Figure 3A) to allow for reaching the target preventive intracellular TFV-DP PBMC concentrations of 100.00 fmol/10^6^ cells (Figure 2A). Animals in both PrEP (n=8) and control (n=6) cohorts were rectally challenged weekly with low-dose SHIV_SF162P3_ for up to 10 inoculations and continually monitored for drug PK throughout the study (Figure 3A). The SHIV inoculation dosage used are similar to human semen HIV RNA levels during acute viremia, thus recapitulating high-risk or acute HIV infection in humans. Therefore, this animal model is considered more aggressive, as the risk of infection per exposure markedly exceeds the risk in clinical settings.^[31]^

**Figure 3.**
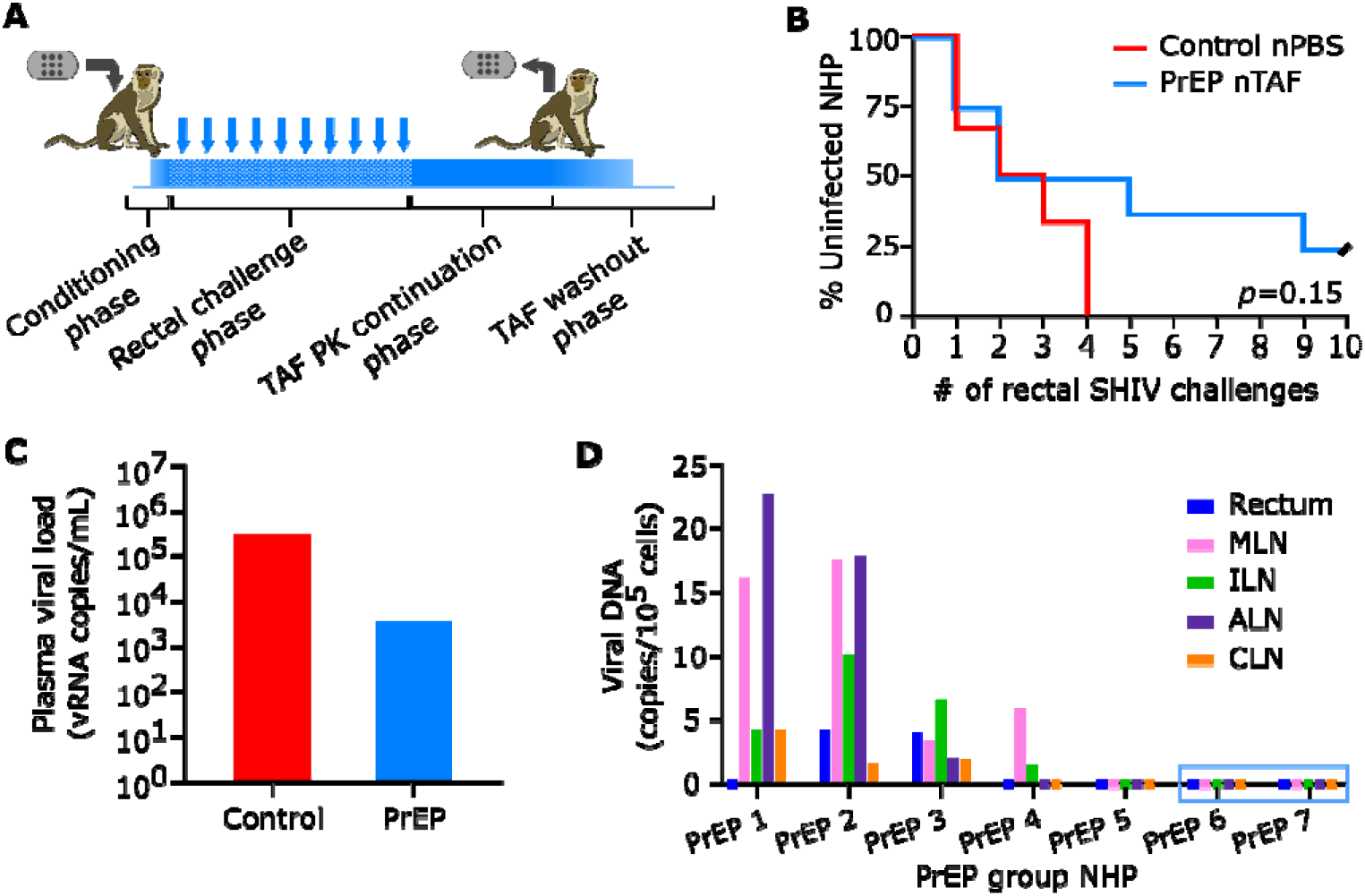
PrEP efficacy of nTAF. A) Schematic of study design. Conditioning phase to reach TFV-DP PBMC concentrations above 100 fmol/10^6^ cells. Rectal challenge phase with up to 10 weekly low-dose SHIV_SF162P3_ exposures. TAF PK continuation phase followed by nTAF explantation from all animals and euthanasia of 4 animals. TAF washout was observed in the remaining 3 animals for 2 months prior to euthanasia. B) Kaplan-Meier curve representing the percentage of infected animals as a function of weekly SHIV exposure. PrEP (n=8) vs control (n=6) group; censored animals represented with black slash. Statistical analysis by Mantel-Cox test. C) Median peak viremia levels in breakthrough animals at initial viral load detection. D) Cell-associated viral DNA loads of tissues in PrEP group. Animals PrEP 1-5 were infected while PrEP 6 and 7 (blue box) remained uninfected throughout the study. MLN, mesenteric lymph nodes, ILN, inguinal lymph nodes, ALN, axillary lymph nodes, CLN, cervical lymph nodes.

To monitor for SHIV_SF162P3_ infection, we evaluated weekly cell-free viral RNA in the plasma. Rectal challenges were stopped upon initial detection of plasma viral RNA, which was confirmed after a consecutive positive assay. Two of eight macaques from the nTAF group (25.00%) were uninfected after 10 weekly rectal SHIV_SF162P3_ challenges (Figure 3B). Based on the number of infections per total number of challenges, the nTAF group had a reduced risk of infection per-exposure of 62.50% (95% CI, 1.72% to 85.69%; *p*=0.068), in comparison to the control group. However, because of the small sample size, the result is not very precise, as indicated by the lower bound of the confidence interval. Prophylaxis with nTAF increased the median time to infection to 5 challenges compared to 2 challenges in the control cohort (*p*=0.38). After device explantation, there was no spike in viremia, indicative of PrEP efficacy of nTAF monotherapy in the two uninfected animals. While Kaplan-Meier analysis demonstrated delayed and reduced infection in some animals, there was no statistical significance (*p*=0.15) between nTAF and nPBS groups.

TAF-treated infected NHPs had blunted SHIV RNA peak viremia (median; 3.80 × 10^4^ vRNA copies/mL; IQR, 1.60 × 10^3^ to 2.09 ×10^5^ vRNA copies/mL) in comparison to control groups (median; 3.01 × 10^5^ vRNA copies/mL; IQR, 9.00 × 10^3^ to 7.25 × 10^6^ vRNA copies/mL) (Figure 3C). However, differences in SHIV RNA levels at initial detection were not statistically significant between control and infected PrEP animals (*p*=0.18 by Mann-Whitney test).

At euthanasia, we assessed the residual SHIV infection in various tissues collected from the nTAF cohort by measuring cell-associated SHIV_SF162P3_ provirus DNA (Figure 3D). Tissues from PrEP 1-4 were assessed after 4 months of nTAF implantation, and after 2 months of drug washout for PrEP 5-7. SHIV DNA was detectable in the MLN in 4/5 of the infected PrEP NHPs. Animals PrEP 5 (infected) and PrEP 6 and 7 (uninfected), had no detectable SHIV DNA in any of the tissues analyzed.

### 2.4. Drug stability in vivo within nTAF

To evaluate drug stability in nTAF after 4 months of *in vivo* implantation, we extracted residual contents from the implant and analyzed for TAF and its hydrolysis products (TAF*) (Table 2). Residual drug within the implant ranged 30.75 – 71.12% of the initial loaded amount. Further, TAF* within the implant was predominantly composed of TAF hydrolysis products, including TFV, with TAF stability ranging 18.21 - 43.08%. Therefore, augmented TAF hydrolysis to TFV within the implant most likely contributed to increased TFV levels observed in plasma towards the end of the study. The nTAF implants had a mean release rate of 1.40 ± 0.39 mg/day, which was sufficient to sustain intracellular TFV-DP concentrations above 100.00 fmol/10^6^ PBMCs throughout the duration of the study.

**Table 2.** Residual drug analysis from nTAF implants at explantation via high performance liquid chromatography (HPLC) and UV-Vis spectroscopy.

| NHP<br>PrEP | TAF loaded<br>(mg) | Residual TAF* (mg) | TAF<br>stability (%) | TAF release rate<br>(mg/day) |
| --- | --- | --- | --- | --- |
| 1 | 341.50 | 161.87 | 30.76 | 1.60 |
| 2 | 330.10 | 217.65 | 12.28 | 1.00 |
| 3 | 337.10 | 215.57 | 18.21 | 1.09 |
| 4 | 382.10 | 241.01 | 31.78 | 1.26 |
| 5 | 457.60 | 325.43 | 43.08 | 1.18 |
| 6 | 449.30 | 279.46 | 18.70 | 1.52 |
| 7 | 342.60 | 105.34 | 22.26 | 2.12 |

### 2.5. nTAF safety and tolerability in NHP

To assess nTAF safety and tolerability, we histologically examined the tissue surrounding the implants after 4 months of implantation, through immunohistochemical analysis (Figure 4A). Specifically, we evaluated the fibrotic capsule in contact with either the titanium reservoir (Figure 4B) or TAF-eluting nanofluidic membrane (Figure 4C and D). Histological analysis via hematoxylin and eosin (H&E) demonstrated foreign-body response, which is typical of medical implants. The surrounding subcutaneous tissue and underlying skeletal muscle was healthy with limited necrosis in the fibrotic capsule. While fibrotic capsules exhibited cellular infiltration, they were negative for inflammatory cell marker CD45 (Figure 4E). DAPI staining demonstrated healthy nuclei in the areas with increased cellular infiltration. Further, analysis of the fibrotic area in contact with TAF-releasing membrane via acid-fast bacteria (AFB) (Figure 4F) and Grocott methenamine silver staining (Figure 4G), which evaluates for presence of bacteria and fungi, respectively, were negative.

**Figure 4.**
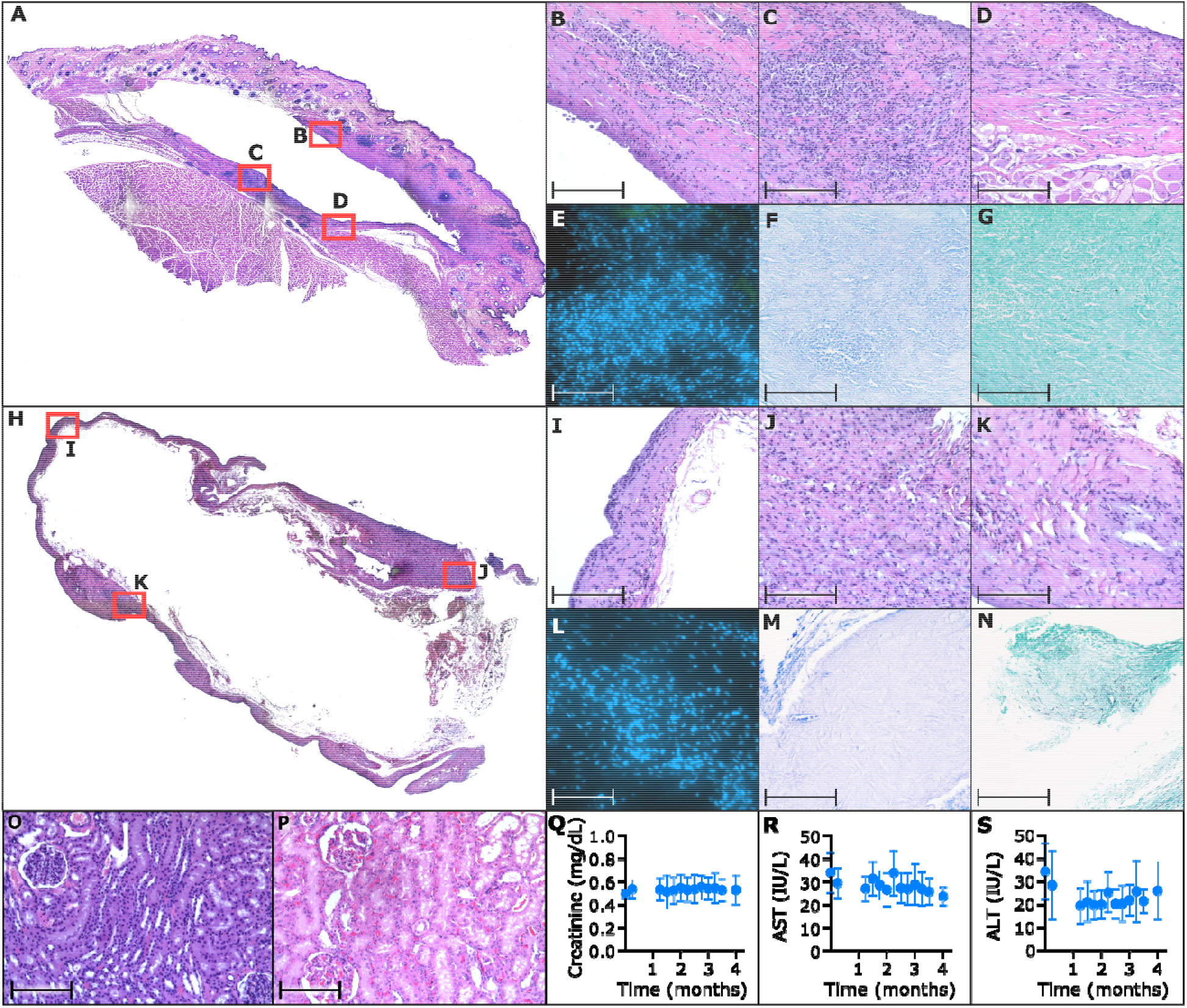
Histological inflammatory response to nTAF and control nPBS; and toxicology assessment of nTAF in the kidney and liver. A) Representative H&E stain of NHP skin surrounding PrEP nTAF, with B) fibrotic capsule in contact with titanium implant; 20 × magnification. Fibrotic capsule in contact with TAF-releasing membrane was assessed via C), D) H&E, 20 × magnification; E) immunofluorescence staining of CD45 (green) and DAPI nuclear stain (blue), 100 × magnification; F) AFB staining for presence of bacteria, 20 × magnification; and G) Grocott methenamine silver staining for presence of fungi, 20 × magnification. H) Representative H&E stain of NHP skin surrounding control nPBS. Fibrotic capsule in contact with titanium implant was assessed via I), J), K) H&E, 20 × magnification, L) immunofluorescence staining of CD45 (green) and DAPI nuclear stain (blue), 100 × magnification; M) AFB staining for presence of bacteria, 20 × magnification; and N) Grocott methenamine silver staining for presence of fungi, 20 × magnification. O) Representative H&E stain of kidney from PrEP nTAF group demonstrating normal histology, in comparison to P) representative H&E stain of kidney from control NHP similarly showing no nephrotoxicity; 20 × magnification. Q) Creatinine activity measurements from nTAF cohort. Liver enzymes, R) aspartate aminotransferase (AST), and S) alanine aminotransferase (ALT) from nTAF cohort. Baseline levels (0 month) were measured before implantation of nTAF. All data are presented as mean ± SD (n=7). Images A and H taken at 4 × magnification and stitched together. Scale bar in 20 and 100 × magnification is 200 and 10 μm, respectively.

In parallel, as a control, the tissue surrounding nPBS implants were histologically assessed (Figure 4H), specifically the fibrotic capsule (Figure 4I-K), which was thinner and denser than the nTAF. Similarly, the tissue surrounding the control implant was negative for CD45 cells (Figure 4L), bacteria (Figure 4M) or fungi (Figure 4N). While other groups have reported that TAF induced necrosis at sites of implantation ^[10]^, overall our results showed no cellular damage or aberrant inflammatory cell influx, indicative of implant tolerability.

As TFV is implicated in nephrotoxicity and hepatotoxicity, we evaluated the kidney and liver in the animals with nTAF implants. The kidney of an untreated NHP from a prior study was used as a historical control, because nPBS NHPs were transferred to another study after infection. Histological assessment of the kidney from nTAF cohort via H&E analysis (Figure 4O) did not demonstrate necrosis or signs of damage, in comparison to control (Figure 4P). Further, creatinine levels were within normal limits throughout the study, suggesting that there was no detectable kidney damage in the nTAF cohort (Figure 4Q). Liver enzymes were monitored as surrogate markers for health; aspartate aminotransferase (AST) (Figure 4R), and alanine aminotransferase (ALT) (Figure 4S) measurements were within normal levels with respect to baseline values pre-nTAF implantation. Metabolic panel, complete blood count and urinalysis results were also within normal levels (Figure S3A-V, 4A-N, Table S1, Supporting Information).

## 3. Discussion

This work represents the first ever preventive efficacy assessment of an implantable LA ARV platform and the foremost study of LA TAF as a single agent HIV PrEP regimen. Our finding that nTAF protected from SHIV infection with 62.50% reduction in risk of infection per exposure resembles that of TAF predecessor, tenofovir disoproxil fumarate (TDF). TDF monotherapy resulted in 60.00% protective efficacy in macaques^[32]^, but clinically achieved 67% risk reduction and 86.00% preventive efficacy in individuals with detectable plasma tenofovir.^[5,33]^

There is no benchmark preventive level of TFV-DP in PBMCs for sustained subcutaneous administration of TAF. We used as a reference the TFV-DP concentration in PBMCs of 100 fmol/10^6^ cells, which conservatively exceeds the levels identified as protective in the iPrEX trial with Truvada® (cryopreserved PBMC, 16.00 fmol/10^6^ cells; freshly lysed PBMC, 40.00 fmol/10^6^ cells).^[6]^ Other TAF-releasing implants are targeting 24-48 fmol/10^6^, a target that takes into consideration the 66% TFV-DP loss during cryopreservation in the iPrEX trial.^[7,9,10]^ While not directly comparable to oral Truvada administration, we used 100 fmol/10^6^ cells as rational target to exceed prior to start the viral challenges. Nonetheless, this is the first efficacy study with continuous TAF administration via the subcutaneous route. Our results show that by maintaining a median TFV-DP concentration of 390 fmol/10^6^ PBMC (IQR, 216.50 to 585.50 fmol/10^6^ PBMC) we achieved partial protection with 62.50% efficacy (95% CI, 1.72% to 85.69%). In light of our studies, it remains unclear what the preventive benchmark could be to establish 100% efficacy in a rectal challenge model.

Most clinical studies evaluating PrEP adherence use plasma, PBMC or dried blood spots as surrogate markers to local tissue concentrations.^[5,6,33,34]^ However, breakthrough infection has occurred in individuals with high systemic drug concentrations, similar to the infected nTAF animals in our study. Therefore, it remains unclear if infection in some animals in our study could be attributable to inadequate TFV-DP concentrations in the site of viral transmission. In a study of weekly oral TAF as a single-agent PrEP against vaginal SHIV infection by the Center for Disease Control, TFV-DP PBMC levels were similar between the four infected and five uninfected animals.^[30]^ However, only five out of nine animals had detectable vaginal TFV-DP concentrations (5 fmol/mg) prior to challenge.^[30]^ It is also of interest to identify the turn-over rate of “TFV-DP positive” to “TFV-DP naïve” mononuclear cells systemically and locally at the site of transmission to improve dosing regimens. Garcia-Lerma et. al demonstrated that once weekly oral TAF dosing conferred low protection from HIV transmission, despite high systemic (>1000 fmol/10^6^ PBMC) and rectal (median, 377 fmol/10^6^ mononuclear cells) TFV-DP levels.^[35]^ However, in the aforementioned study the animals were rectally challenged 3 days after the first weekly oral TAF dose. Thus, the long interval between drug dosing and virus exposure could have allowed for TFV-DP naïve mononuclear cells to repopulate at the site of transmission. Of relevance, on-demand local TFV delivery at HIV transmission sites, such as a TFV rectal douche, has shown to achieve high local tissue concentrations and favorable PK profiles in NHP with SHIV challenges.^[36, 37]^ Therefore, we posit that PrEP efficacy could plausibly be improved if first-line target cells have sufficient TFV-DP concentrations prior to virus exposure.

The present study was limited by the number of animals and the use of both sexes for rectal SHIV prevention. Future studies could address this issue by increasing the sample size and conducting separate sexes studies to evaluate protection against rectal or vaginal exposure. Further, because Descovy® is clinically approved for oral administration, scientific rigor could be strengthened with an additional group with daily oral TAF dosing as opposed to weekly dosing as performed in literature, in comparison to sustained subcutaneous delivery.

In summary, our innovative strategy of continuous low-dose systemic delivery of TAF obviates adherence challenges and provides similar protective benefit to that observed with oral TDF. Taken together, this work provides optimism for implementing clinical studies to assess the safety and efficacy of LA TAF platforms for HIV PrEP.

## 4. Experimental Section

### Nanofluidic implant assembly

Medical-grade 6AI4V titanium oval drug reservoirs were specifically designed and manufactured for this study. Briefly, a nanofluidic membrane possessing 278,600 nanochannels (mean; 194 nm) was mounted on the inside of the sterile drug reservoir as described previously.^[12]^ Detailed information regarding the membrane structure and fabrication was described previously.^[26, 38]^Implants were welded together using Arc welding. PrEP implants were loaded with ~300 - 457 mg TAF fumarate using a funnel in the loading port, while control implants were left empty. A titanium piece that resembled a small nail was inserted into the loading port and welded shut. Implants were primed for drug release through the nanofluidic membrane by placing implants in 1 X Phosphate Buffered Saline (PBS) under vacuum. This preparation method resulted in loading of control implants with PBS. Implants were maintained in sterile 1X PBS in a hermetically sealed container until implantation shortly after preparation. TAF was kindly provided by Gilead Sciences, Inc.

### In vitro release from nanofluidic implant

In an effort to limit the amount of drug used, the in vitro release study was performed using nanochannel membranes with identical structure and channel size those adopted in vivo, but with a small number of nanochannels (n= 9,800 as compared to n=278,600 for the full-size membrane). In vitro release results were then linearly scaled to account for the difference in nanochannels number. Medical-grade 6AI4V titanium cylindrical drug reservoirs (n=5) were assembled as described above, loaded with ~20.00 mg TAF fumarate and placed in sink solution of 20 mL 1 × PBS with constant agitation at 37°C. For analysis, the entire sink solution was retrieved and replaced with fresh PBS every other day for 20 days. The maximum TAF concentration regarding TAF saturation in sink solution was <10%, therefore maintaining sink condition. High-performance liquid chromatography (HPLC) analysis was performed on an Agilent Infinity 1260 system equipped with a diode array and evaporative light scattering detectors using a 3.5-μm 4.6 × 100 mm Eclipse Plus C18 column and water/methanol as the eluent and 25 μL injection volume. Peak areas were analyzed at 260 nm absorbance.

### Nanofluidic membrane assessment

Silicon nanofluidic membranes structure and composition was assessed using different imaging techniques at the Microscopy – SEM/AFM core of the Houston Methodist Research Institute (HMRI), Houston, TX, USA. Inspection of structural conformation was performed via scanning electron microscopy (SEM; Nova NanoSEM 230, FEI, Oregon, USA), nanochannel dimension was measured on membrane cross sections obtained using gallium ion milling (FIB, FEI 235). Surface roughness was measured by atomic force microscopy (AFM Catalyst), surface chemical composition was evaluated with Energy-dispersive X-ray spectroscopy (EDAX, Nova NanoSEM 230).

### Animals and animal care

All animal procedures were conducted at the AAALAC-I accredited Michale E. Keeling Center for Comparative Medicine and Research, The University of Texas MD Anderson Cancer Center (UTMDACC), Bastrop, TX. All animal experiments were carried out according to the provisions of the Animal Welfare Act, PHS Animal Welfare Policy, and the principles of the NIH Guide for the Care and Use of Laboratory Animals. All procedures were approved by the Institutional Animal Care and Use Committee (IACUC) at UTMDACC, which has an Animal Welfare Assurance on file with the Office of Laboratory Animal Welfare. IACUC #00001749-RN00. Indian rhesus macaques (*Macaca mulatta*; n=14; 6 males and 8 females) of 2-4 years and 2-5 kg bred at this facility were used in the study. All procedures were performed under anesthesia with ketamine (10 mg/kg, intramuscular) and phenytoin/pentobarbital (1 mL/10 lbs, intravenous [IV]).

All animals had access to clean, fresh water at all times and a standard laboratory diet. Prior to the initiation of virus inoculations, compatible macaques were pair-housed. Once inoculations were initiated, the macaques were separated into single housing (while permitting eye contact) to prevent the possibility of SHIV transmission between the macaques. Euthanasia of the macaques was accomplished in a humane manner (IV pentobarbital) by techniques recommended by the American Veterinary Medical Association Guidelines on Euthanasia. The senior medical veterinarian verified successful euthanasia by the lack of a heartbeat and respiration.

### Minimally invasive implantation procedure

An approximately 1-cm dorsal skin incision was made on the right lateral side of the thoracic spine. Blunt dissection was used to make a subcutaneous pocket ventrally about 5 cm deep. The implant was placed into the pocket with the membrane facing the body. A simple interrupted tacking suture of 4-0 polydioxanone (PDS) was placed in the subcutaneous tissue to help close the dead space and continued intradermally to close the skin. All animals received a single 50,000 U/kg perioperative penicillin G benzathine/penicillin G procaine (Combi-Pen) injection and subcutaneous once-daily meloxicam (0.2 mg/kg on day 1 and 0.1 mg/kg on days 2 and 3) for postsurgical pain.

### Blood collection and plasma and PBMC sample preparation

All animals had weekly blood draws to assess plasma TAF and TFV concentrations, intracellular TFV-DP PBMC concentrations, plasma viral RNA loads, and cell-associated SHIV DNA in PBMCs. Blood collection and sample preparation were performed as previously described.^[11]^ Blood was collected in EDTA-coated vacutainer tubes before implantation; on days 1, 2, 3, 7, 10, and 14; and then once weekly until euthanasia. Plasma was separated from blood by centrifugation at 1200 × *g* for 10 min at 4 °C and stored at −80 °C until analysis. The remaining blood was used for PBMC separation by standard Ficoll-Hypaque centrifugation. Cell viability was > 95%. After cells were counted, they were pelleted by centrifugation at 400 × *g* for 10 min, resuspended in 500 μL of cold 70% methanol/30% water, and stored at −80 °C until further use.

### Pharmacokinetic analysis of TFV-DP in PBMC and TAF and TFV in plasma

The PK profiles of TFV-DP in PBMC and TAF and TFV in plasma were evaluated throughout the 4 months of nTAF implantation. Due to early implant removal in one animal on day 43, seven animals were evaluated for drug PK. After device explantation, drug washout was assessed for an additional 2 months (n=3).

Intracellular TFV-DP concentrations in PBMCs were quantified using previously described validated liquid chromatographic-tandem mass spectrometric (LC-MS/MS) analysis.^[6, 39]^ The assay was linear from 5 to 6000 fmol/sample. Typically, 25 fmol/sample was used as the lower limit of quantitation (LLOQ). If additional sensitivity was needed, standards and quality controls were added down to 5 fmol/samples, as previously described.^[39]^ Day 21 TFV-DP concentrations were omitted due to PBMC count below threshold.

Plasma TAF and TFV concentrations were quantified using a previously described LC-MS/MS assay.^[40]^ Drugs were extracted from 0.1 mL plasma via solid phase extraction; assay lower limits of quantitation for TAF and TFV were 0.03 ng/mL and 1 ng/mL, respectively. The multiplexed assay was validated in accordance with FDA, Guidance for Industry: Bioanalytical Method Validation recommendations.^[41]^

### Tissue TFV-DP quantification

Lymphoid tissues (mesenteric, axillary, and inguinal lymph nodes), rectum, urethra, cervix, tonsil, spleen, and liver were homogenized, and 50- to 75-mg aliquots were used for TFV-DP quantitation. Pharmacokinetic analysis of TFV-DP was conducted by the Clinical Pharmacology Analytical Laboratory at the Johns Hopkins University School of Medicine. TFV concentrations in aforementioned tissue biopsies were determined via LC-MS/MS analysis. TFV-DP was measured using a previously described indirect approach, in which TFV was quantitated following isolation of TFV-DP from homogenized tissue lysates and enzymatic conversion to the TFV molecule.^[39]^ The assay LLOQ for TFV-DP in tissue was 5 fmol/sample, and drug concentrations were normalized to the amount of tissue analyzed.^[42]^ The TFV-DP tissue was validated in luminal tissue (rectal and vaginal tissue) in accordance with FDA, Guidance for Industry: Bioanalytical Method Validation recommendations^[41]^; alternative tissue types were analyzed using this method.

### PrEP nTAF efficacy against rectal SHIV challenge

To study the efficacy of the PrEP implant against SHIV transmission, animals were divided into two groups, PrEP nTAF-treated [n=8; 4 male (M) and 4 female (F)] or control nPBS (n=6; 3 M and 3 F), in a non-blinded study. The PrEP regimen consisted of subcutaneously implanted nTAF for sustained drug release over 112 days. The efficacy of nTAF in preventing rectal SHIV transmission was evaluated using a repeat low-dose exposure model described previously.^[32,35,43]^ Animals were considered protected if they remained negative for SHIV RNA throughout the study. Briefly, after PrEP-treated macaques achieved intracellular TFV-DP concentrations above 100.00 fmol/10^6^ PBMCs, both groups were rectally exposed to SHIV_SF162P3_ once a week for up to 10 weeks until infection was confirmed by two consecutive positive plasma viral RNA loads. The SHIV_SF162P3_ dose was in range of HIV-1 RNA levels found in human semen during acute viremia.^[43]^

Challenge stocks of SHIV_162p3_ were generously supplied by Dr. Nancy Miller, Division of AIDS, NIAID, through Quality Biological (QBI), under Contract No. HHSN272201100023C to the Vaccine Research Program, Division of AIDS, NIAID. The stock SHIV_162p3_ R922 derived harvest 4 dated 9/16/2016 (p27 content 173.33 ng/ml, viral RNA load >10^9^ copies/ml, TCID50/ml in rhesus PBMC 1280) was diluted 1:300 and 1ml of virus was used for rectal challenge each time.

For the challenge, the animals were positioned in prone position and virus was inoculated approximately 4 cm into the rectum. Inoculated animals were maintained in the prone position with the perineum elevated for 20 minutes to ensure that virus did not leak out. Care was also taken to prevent any virus from contacting the vagina area and to not abrade the mucosal surface of the rectum.

### Infection monitoring by SHIV RNA in plasma and SHIV DNA in tissues

Infection was monitored by the detection of SHIV RNA in plasma using previously described methods^[44, 45]^ with modification. Viral RNA (vRNA) was isolated from blood plasma using the Qiagen QIAmp UltraSense Virus Kit (Qiagen #53704) in accordance with manufacturer’s instructions for 0.5 mL of plasma. vRNA levels were determined by quantitative real-time PCR (qRT-PCR) using Applied Biosystems^TM^ TaqMan^TM^ Fast Virus 1-Step Master Mix (Thermofisher #4444432) and a primer-probe combination recognizing a conserved region of gag (GAG5f: 5’-ACTTTCGGTCTTAGCTCCATTAGTG-3’; GAG3r: 5’-TTTTGCTTCCTCAGTGTGTTTCA-3’; and GAG1tq: FAM 5’-TTCTCTTCTGCGTGAATGCACCAGATGA-3’TAMRA). Each 20 μl reaction contained 900 nM of each primer and 250 nM of probe, and 1x Fast Virus 1-Step Master Mix, plasma-derived vRNA sample, SIV gag RNA transcript containing standard, or no template control.

qRT-PCR was performed in a ABI Step One Plus Cycler. PCR was performed with an initial step at 50°C for 5 min followed by a second step at 95°C for 20 sec, and then 40 cycles of 95°C for 15 sec and 60°C for 1 min. Ten-fold serial dilutions (1 to 1 × 10^6^ copies per reaction) of an in vitro transcribed SIV gag RNA were used to generate standard curves. Each sample was tested in duplicate reactions. Plasma viral loads were calculated and shown as viral RNA copies/mL plasma. The limit of detection is 50 copies/ml. Infections were confirmed after a consecutive positive plasma viral load measurement.

To detect viral DNA in tissue samples, total DNA was isolated from PBMCs or tissue specimens using the Qiagen DNeasy Blood & Tissue Kit (Qiagen #69504) according to the manufacturer’s protocol. DNA was quantified using a nanodrop spectrophotometer. qRT-PCR was performed using the SIV gag primer probe set described above. Each 20 μl reaction contained 900 nM of each primer and 250 nM of probe, and 1x TaqMan Gene Expression Master Mix (Applied Biosystems, Foster City, CA), macaque-derived DNA sample, SIV gag DNA containing standard, or no template control. PCR was initiated in with an initial step of 50°C for 2 min and then 95°C for 10 min. This was followed by 40 cycles of 95°C for 15 sec, and 60°C for 1 min. Each sample was tested in triplicate reactions. Ten-fold serial dilutions of a SIV gag DNA template (1 to 1 × 10^5^ per reaction) were used to generate standard curves. The limit of detection of this assay was determined to be 1 copy of SIV gag DNA.

### Device retrieval and macaque euthanasia

A subset of PrEP-treated macaques (n=4), those with the highest viral load, were euthanized on day 112, while implants were retrieved on day 112 from the remaining PrEP-treated macaques (n=3) for continuation to a 2-month drug-washout period before euthanasia. SHIV-infected macaques in the control group (n=6) were transferred to another study (data not shown) and euthanized 28 days later. The implant was retrieved with a small incision in the skin and stored at −80 °C until further analysis. Skin within a 2-cm margin surrounding the implant was excised from euthanized macaques and fixed in 10% buffered formalin for histological analysis. Macaques continuing in the washout period underwent a skin punch biopsy of the subcutaneous pocket, and the skin incision was sutured with a simple interrupted tacking suture of 4-0 PDS; the specimen was fixed in 10% buffered formalin for histological analysis. The following tissues were collected from all animals at euthanasia (n=13): lymphoid tissues (mesenteric, axillary, and inguinal lymph nodes), rectum, urethra, cervix, tonsil, spleen, and liver. Tissues were snap-frozen and stored at −80 °C until further analysis of TAF concentrations, viral RNA loads, and cell-associated SHIV DNA.

### Residual drug and nanofluidic membrane retrieval from explanted implants

Upon explantation, the implants were snap frozen with liquid nitrogen to preserve residual drug for stability analysis. For residual drug retrieval, the implants were thawed at 4°C overnight. A hole was drilled on the outermost corner on the back of the implant using a 3/64 titanium drill bit with a stopper. Drilling was performed on the back of the implant and distal to the membrane to avoid damage. Following drilling, 20 μL sample from the implant drug reservoir was aliquoted into respective 1.5 mL Eppendorf tubes with 0.5 mL 100% ethanol using a pipette. The implants were placed in 50 mL conical tubes with 40.0 g 70% ethanol. Each implant was flushed using a 19-gauge needle with 70% ethanol from the sink solution. For sterilization, the implants were incubated in 70% ethanol for 4 days and transferred to new conical tubes with fresh 70% ethanol for an additional 4 days. To ensure nanochannel membranes were dry, the implants were transferred to new conical tubes with 100% ethanol for a day and placed in 6-well plates to dry under vacuum. To protect the membrane during machining procedure, electrical tape was placed over the outlets. The implants were opened using a rotary tool with a diamond wheel. Titanium dust from machining procedure was gently cleaned from membrane with a cotton swab and 70% ethanol. To remove membrane from the implant, a drop of nitric acid (Trace Metal grade) was placed on the membrane overnight and rinsed with Millipore water the next day. Membranes were kept in hermetically sealed containers until analysis.

### TAF stability analysis in drug reservoir

Liquid in the drug reservoir after explantation was collected with a pipette and diluted 25 times with 100% ethanol. The samples were transferred to 0.2 μm nylon centrifugal filters and centrifuged at 500 G for 8 minutes at room temperature. An aliquot of 50 μL from the filtered samples were further diluted in 100 μL 100% ethanol. HPLC analysis was performed on an Agilent Infinity 1260 system equipped with a diode array and evaporative light scattering detectors using a 3.5-μm 4.6 × 100 mm Eclipse Plus C18 column and water/methanol as the eluent and 25 μL injection volume. Peak areas were analyzed at 260 nm absorbance.

Drug solids from within the implant were analyzed from the initial 40.0 g 70% ethanol sink solution. The samples were transferred to 0.2 μm nylon centrifugal filter and centrifuged at 500 G for 8 minutes at room temperature. An aliquot of 10 μL from the filtered samples was further diluted in 990 μL of deionized water. UV-vis spectroscopy was performed on a Beckman Coulter DU® 730 system. Peak areas were analyzed at 260 nm absorbance.

### Assessment of PrEP nTAF safety and tolerability

Tissues were fixed in 10% buffered formalin and stored in 70% ethanol until analysis. Tissues were then embedded in paraffin, cut into 5 μm sections and stained with hematoxylin and eosin (H&E) staining at the Research Pathology Core HMRI, Houston, TX, USA. H&E staining was performed on tissue sections surrounding the implant site and kidney. Histological assessment was performed by a blinded pathologist. For immunohistochemistry evaluation of tissue sections, slides were stained with anti-CD45 conjugated to fluorescein isothiocyanate (Pharmingen). For negative controls, corresponding immunoglobulin and species (IgG)-matched isotype control antibodies were used. Nonspecific binding in sections was blocked by a 1-hour treatment in tris-buffered saline (TBS) plus 0.1% w/v Tween containing defatted milk powder (30 mg ml^−1^). Stained sections were mounted in Slow Fade GOLD with 4’,6-diamidino-2-phenylindole (DAPI) (Molecular Probes, OR) and observed using a Nikon T300 Inverted Fluorescent microscope (Nikon Corp., Melville, NY). For verification of cell phenotype, each slide was scored by counting three replicate measurements by the same observer for each slide. All slides were counted without knowledge of the cell-specific marker being examined, and results were confirmed through a second reading by another observer.

### Assessment of TAF toxicity

To assess TAF toxicity, a comprehensive metabolic panel was analyzed for each animal weekly during the rectal challenge phase of the study and biweekly afterward. Urine and CBCs were analyzed monthly to assess kidney and liver function and monitor the well-being of the NHPs.

### Statistical analysis

Plasma t_1/2_ PK analysis was performed in Microsoft Excel using 2 time points, days 112 and 119. Results were expressed as actual t_1/2_ is less than obtained t_1/2_ (because day 119 values were undetectable and were substituted with BLOQ values). PBMC PK analysis was performed using PKSolver add-in for Microsoft Excel developed by Zhang et al.^[46]^ Data are represented as mean ± SD or median with interquartile range (IQR) between the first (25th percentile) and third (75th percentile) quartiles. The relative risk and relative risk reduction with 95% confidence intervals (95% CI) were estimated to examine the per-exposure effect of TAF, and the Fisher’s exact was used for the comparison. The Mann-Whitney test was used to compare the median survival time and the differences in SHIV RNA levels at initial detection. The Kaplan-Meier analysis was performed between the PrEP and control groups, with the use of the number of inoculations as the time variable. The exact log-rank test was used to test the survival between the two groups. All statistical analysis for calculation of the efficacy of TAF were performed with GraphPad Prism 8 (version 8.2.0; GraphPad Software, Inc., La Jolla, CA). Statistical significance was defined as two-tailed p<0.05 for all tests.

## Supporting Information

Supporting Information is available from the Wiley Online Library or from the author.

## Acknowledgements

We thank Dr. Andreana L. Rivera, Yuelan Ren, and Sandra Steptoe from the research pathology core of Houston Methodist Research Institute. Dr. Jianhua “James” Gu from the electron microscopy core. We thank Simone Capuani from the Houston Methodist Research Institute for implant design and rendering, Dixita Viswanath for help with tissue dissection, and Nicola Di Trani for obtaining FIB image. We thank Dr. Dorothy Lewis for the useful discussions. We thank Luke Segura, Elizabeth Lindemann and Dr. Greg Wilkerson from the Michale E. Keeling Center for Comparative medicine and Research at UTMDACC for support in animal studies and Bharti Nehete for plasma and PBMC isolation and virus challenge preparation. TAF fumarate was provided by Gilead Sciences, Inc.

## Author contributions

**Fernanda P. Pons-Faudoa**: conceptualization, formal analysis, investigation, writing-original draft preparation, writing-review and editing, visualization, project administration. **Antons Sizovs**: conceptualization, methodology, formal analysis, investigation, writing-review and editing, visualization. **Kathryn A. Shelton**: investigation. **Zoha Momin**: investigation. **Lane R. Bushman**: methodology, validation, investigation. **Jiaqiong Xu:**formal analysis. **Corrine Y. X. Chua**: formal analysis, writing-review and editing. **Joan E. Nichols**: methodology, investigation, resources, writing-reviewing and editing. **Trevor Hawkins:**conceptualization, writing-review and editing. **James F. Rooney:**conceptualization, writing-review and editing. **Mark A. Marzinke**: methodology, validation, investigation, resources, data curation, writing-review and editing. **Jason T. Kimata**: validation, investigation, resources, writing-review and editing. **Peter L. Anderson**: methodology, validation, resources, writing-review and editing. **Pramod N. Nehete**: investigation, resources, project administration, writing-review and editing. **Roberto C. Arduino**: conceptualization, writing-review and editing. **Mauro Ferrari:**writing-review and editing. **K. Jagannadha Sastry**: conceptualization, resources, writing-review and editing. **Alessandro Grattoni**: conceptualization, investigation, resources, writing-original draft preparation, writing-review and editing, visualization, supervision, project administration, funding acquisition.

**Correspondence and requests for materials** should be addressed to A.G.

## Table of Contents (ToC)

### ToC text

The Grattoni group performed the first HIV pre-exposure prophylaxis (PrEP) assessment of an implantable long-acting antiretroviral platform. In this foremost study, the partial protection of simian HIV with tenofovir alafenamide (TAF) delivered by a nanofluidic implant was demonstrated in nonhuman primates.

### ToC figure

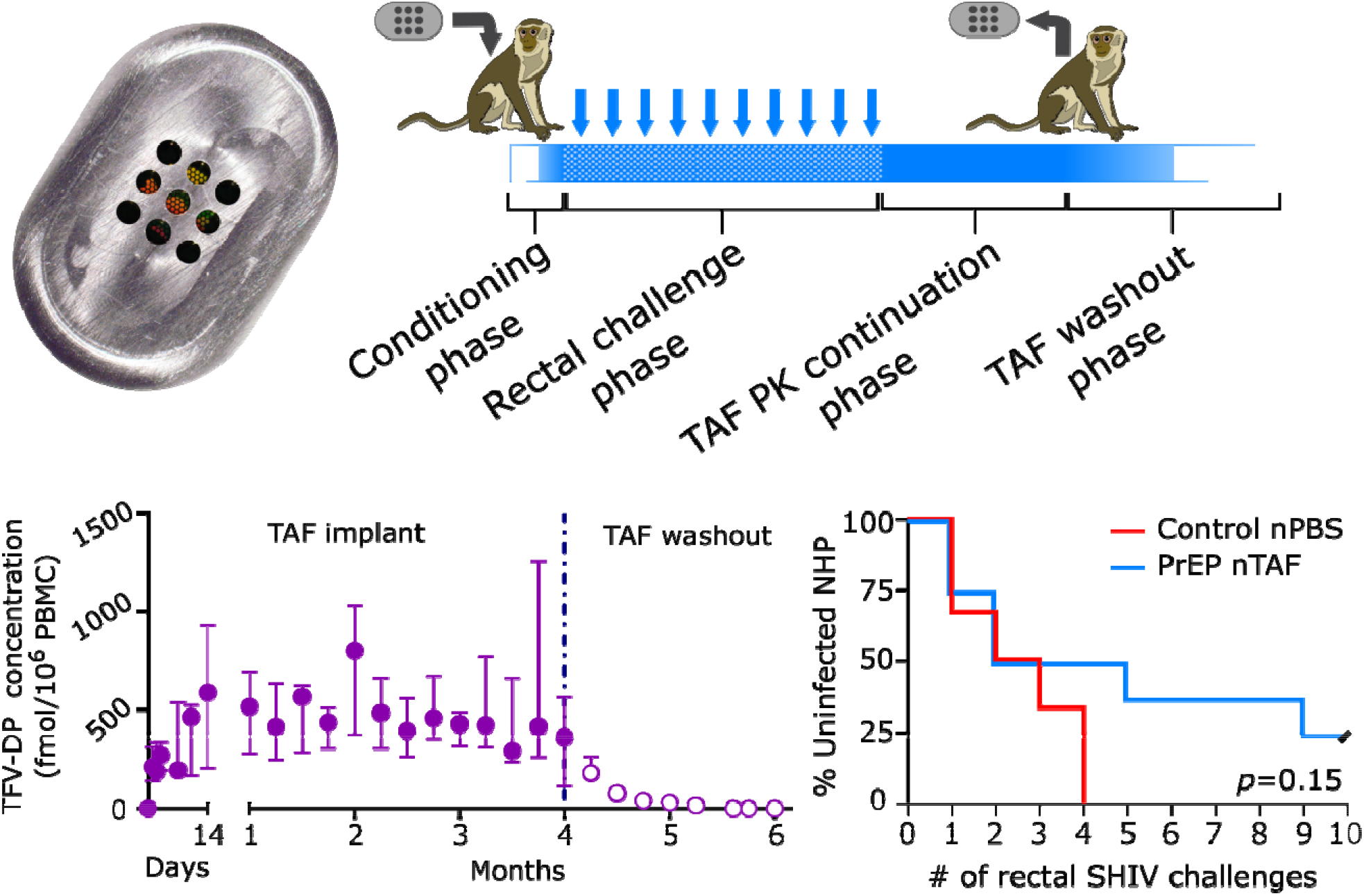

## Supporting Information

**Supporting Figure S1.**
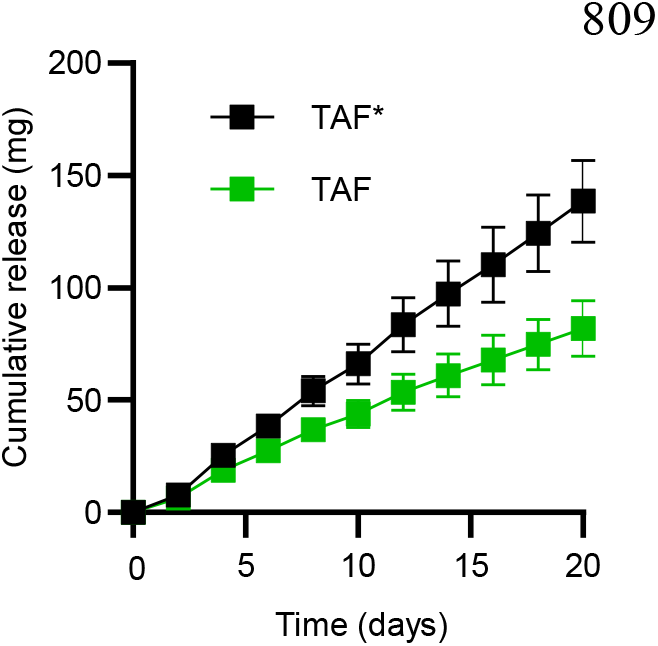
Cumulative in vitro release of TAF from nTAF (n=5). Sum of TAF* shown in black. Data presented as mean ± SEM.

**Supporting Figure S2.**
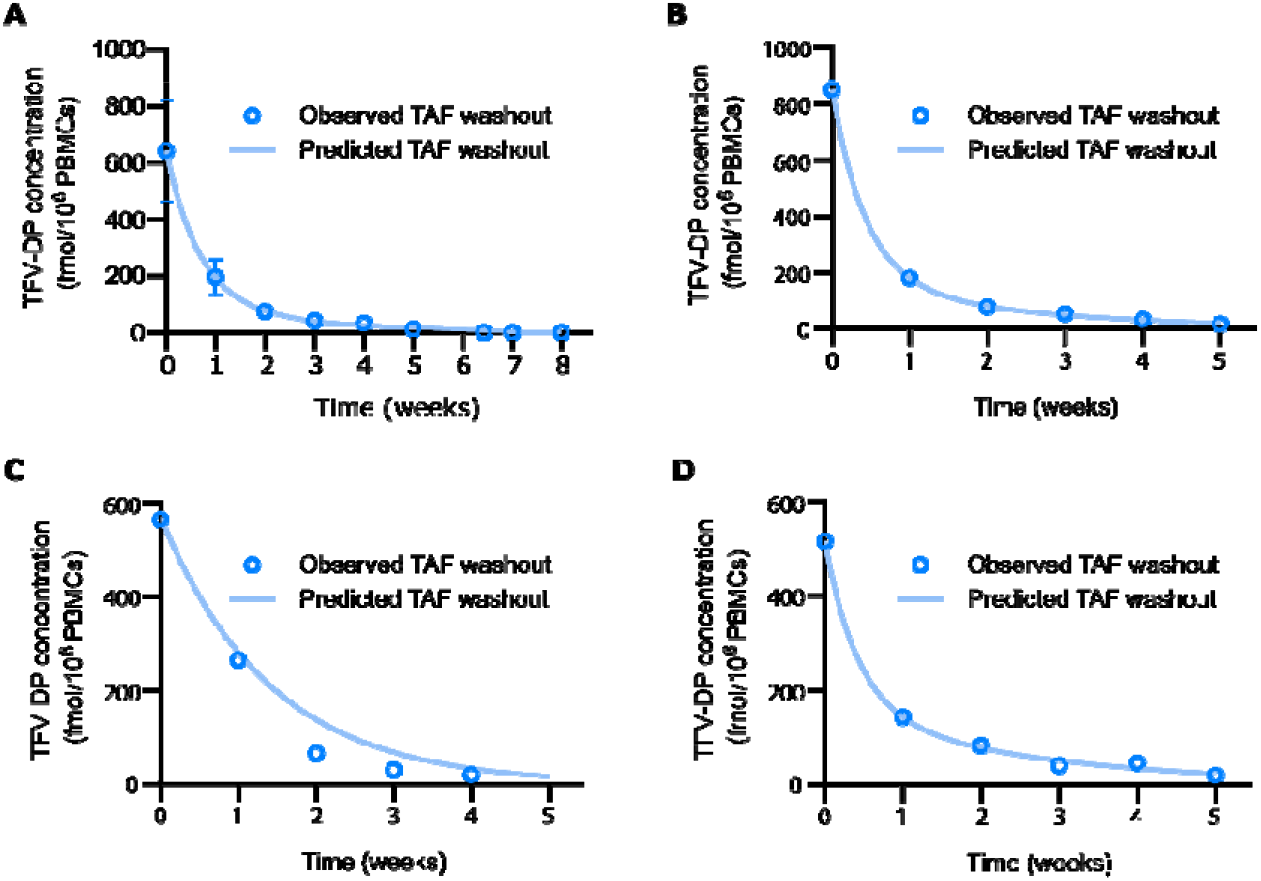
(A) nTAF TFV-DP washout fitted to intravenous bolus injection two-compartment model to determine elimination rate constant. Data are presented as mean ± SD. (B) PrEP 5 nTAF TFV-DP washout fitted to intravenous bolus injection two-compartment mode to determine elimination rate constant. (C) PrEP 6 nTAF TFV-DP washout fitted to intravenous bolus injection two-compartment model to determine elimination rate constant. (D) PrEP 7 nTAF TFV-DP washout fitted to intravenous bolus injection two-compartment model to determine elimination rate constant.

**Supporting Figure S3.**
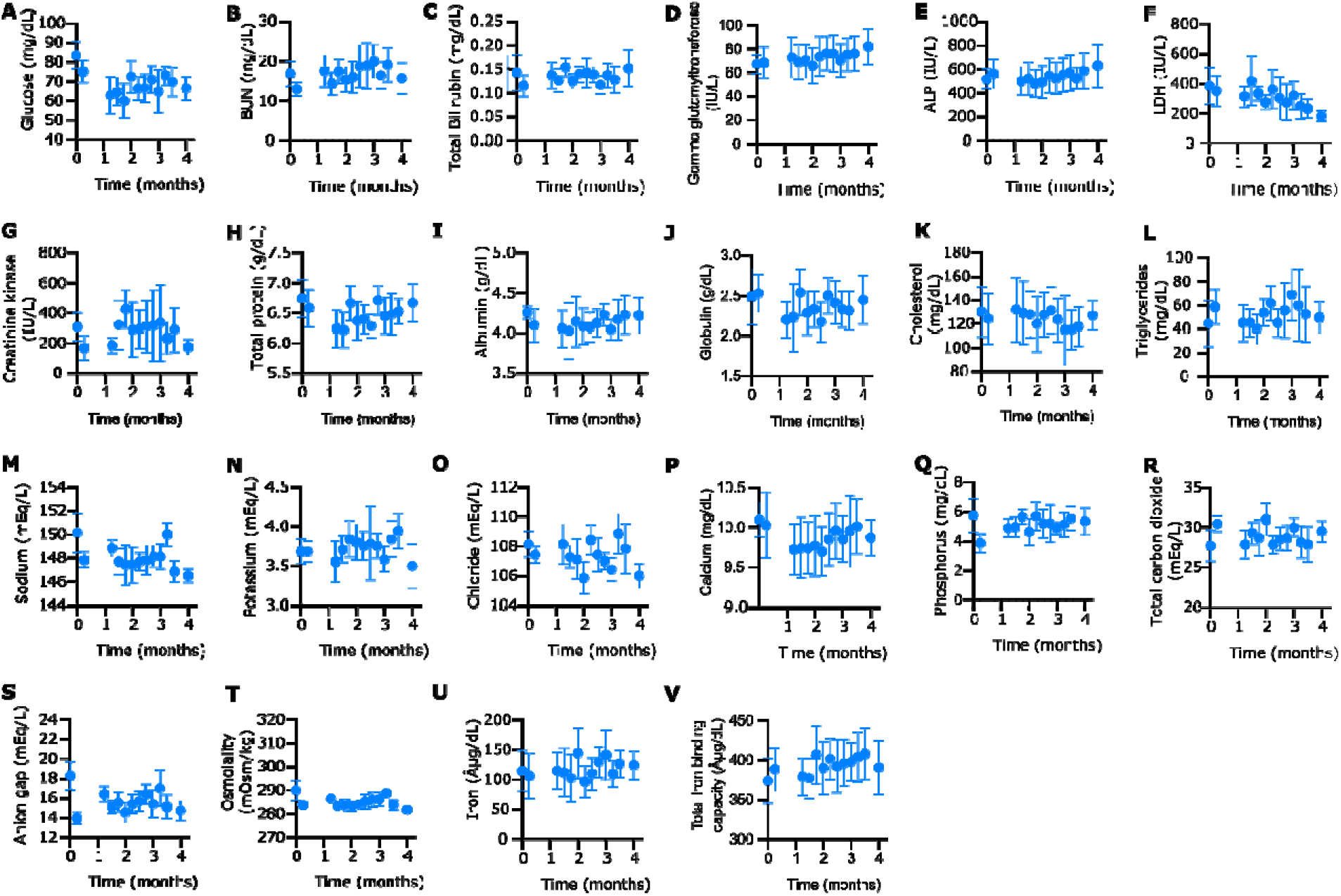
Metabolic panel of rhesus macaques with nTAF. Baseline value for comparison is on day 0 pre-implantation. All data presented as mean ± SD.

**Supporting Figure S4.**
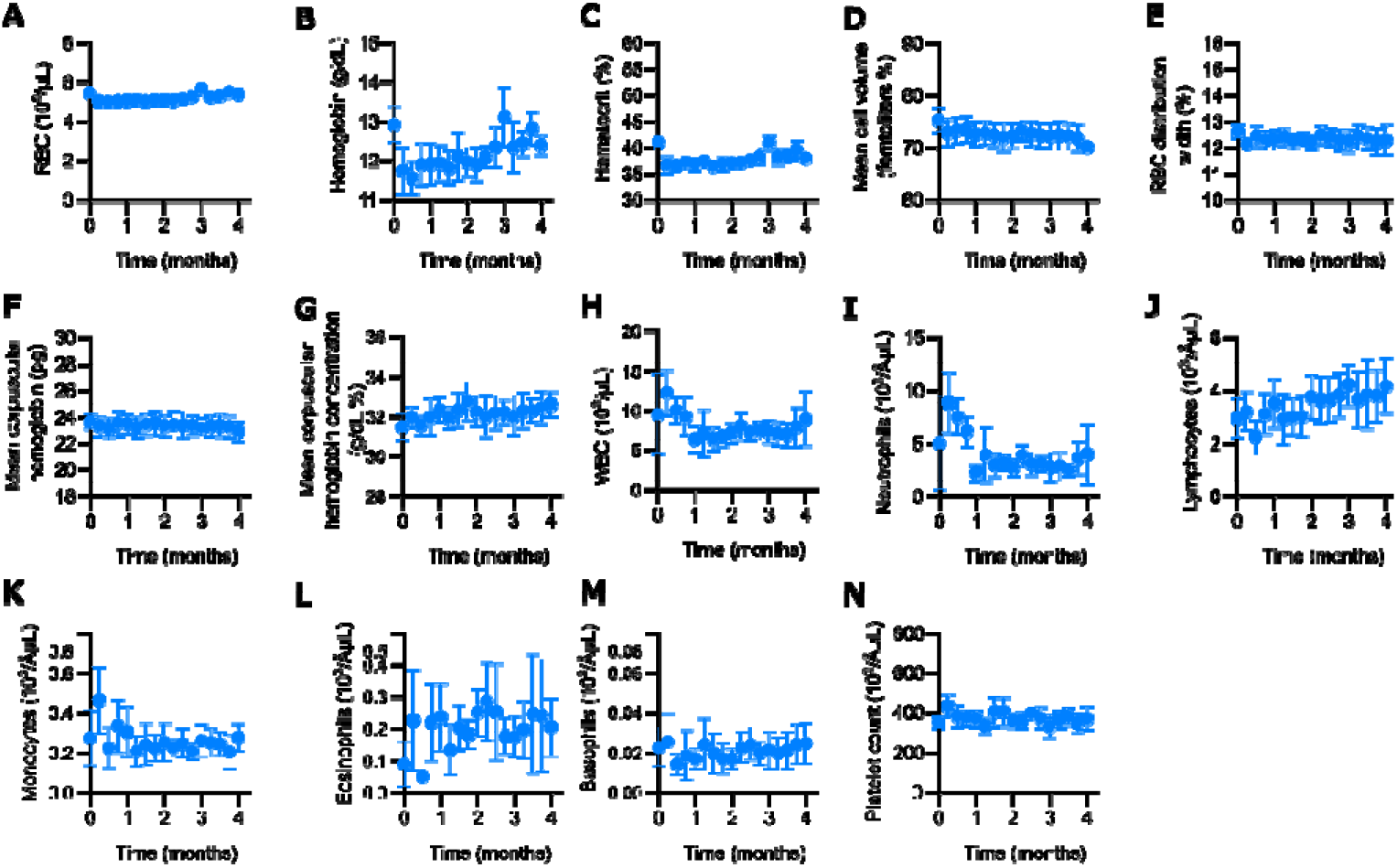
CBC of rhesus macaques with nTAF. Baseline value for comparison is on day 0 pre-implantation. All data presented as mean ± SD.

**Supporting Table S1.** Urinalysis results in rhesus macaques with nTAF. Found in Excel File named: Supporting Table S1.

