## Supplementary Information Table S1 for "Preventive efficacy of a tenofovir alafenamide fumarate nanofluidic implant in SHIV-challenged nonhuman primates"

| Animal | Sex | DOB | Date | Uglu (Mg/Dl) | Ubil () | Uket (Mg/Dl) |
|---|---|---|---|---|---|---|
| PrEP 1 15-166 | M | 16-Sep-15 | 12-Feb-19 | Negative | Negative | Negative |
| | | | 19-Feb-19 | Negative | Negative | Negative |
| | | | 26-Feb-19 | Negative | Negative | Negative |
| | | | 5-Mar-19 | Negative | Negative | Negative |
| | | | 12-Mar-19 | Negative | Negative | Negative |
| PrEP 2 17-087 | F | 16-May-17 | 12-Feb-19 | Negative | Negative | Negative |
| | | | 19-Feb-19 | Negative | Negative | Negative |
| | | | 26-Feb-19 | Negative | Negative | Negative |
| | | | 5-Mar-19 | Negative | Negative | Negative |
| | | | 12-Mar-19 | Negative | Negative | Negative |
| PrEP 3 17-063 | F | 27-Apr-17 | 12-Feb-19 | Negative | Negative | Negative |
| | | | 19-Feb-19 | Negative | Negative | Negative |
| | | | 26-Feb-19 | Negative | Negative | Negative |
| | | | 5-Mar-19 | Negative | Negative | Negative |
| | | | 12-Mar-19 | Negative | Negative | Negative |
| PrEP 4 15-112 | M | 28-Jun-15 | 12-Feb-19 | Negative | Negative | Negative |
| | | | 19-Feb-19 | Negative | Negative | Negative |
| | | | 26-Feb-19 | Negative | Negative | Negative |
| | | | 5-Mar-19 | Negative | Negative | Negative |
| | | | 12-Mar-19 | Negative | Negative | Negative |
| PrEP 5 16-133 | F | 26-Aug-16 | 12-Feb-19 | Negative | Negative | Negative |
| | | | 19-Feb-19 | Negative | Negative | Negative |
| | | | 26-Feb-19 | Negative | Negative | Negative |
| | | | 5-Mar-19 | Negative | Negative | Negative |
| | | | 12-Mar-19 | Negative | Negative | Negative |
| PrEP 6 16-139 | F | 19-Aug-16 | 12-Feb-19 | Negative | Negative | Negative |
| | | | 19-Feb-19 | Negative | Negative | Negative |
| | | | 26-Feb-19 | Negative | Negative | Negative |
| | | | 5-Mar-19 | Negative | Negative | Negative |
| | | | 12-Mar-19 | Negative | Negative | Negative |
| PrEP 7 17-010 | M | 3-Mar-17 | 12-Feb-19 | Negative | Negative | Negative |
| | | | 19-Feb-19 | Negative | Negative | Negative |
| | | | 26-Feb-19 | Negative | Negative | Negative |
| | | | 5-Mar-19 | 100 | Negative | Negative |
| | | | 12-Mar-19 | Negative | Negative | Negative |

| Ubld () | 'Upro (Mg/Dl)' | 'Uro (E.U./Dl)' | 'Nitr ()' | 'Leu ()' | Cm () | Col () | Appe () |
|---|---|---|---|---|---|---|---|
| Negative | Negative | 0.2 | Negative | Negative | - | Yellow | - |
| Trace-int | Negative | 0.2 | Negative | Negative | - | Yellow | - |
| Negative | Negative | 0.2 | Negative | Negative | - | Yellow | - |
| Negative | Negative | 0.2 | Negative | Negative | - | Yellow | - |
| Negative | Negative | 0.2 | Negative | Negative | - | Yellow | - |
| Trace-lys | Negative | 0.2 | Negative | Negative | - | Yellow | - |
| Small | Negative | 0.2 | Negative | Trace | - | Yellow | - |
| Negative | Negative | 0.2 | Positive | Negative | - | Yellow | - |
| Trace-int | Negative | 0.2 | Negative | Negative | - | Straw | - |
| Negative | Negative | 0.2 | Negative | Negative | - | Yellow | - |
| Trace-lys | Negative | 0.2 | Negative | Negative | - | Yellow | - |
| Large | Negative | 0.2 | Negative | Moderate | - | Yellow | - |
| Negative | Negative | 0.2 | Negative | Negative | - | Yellow | - |
| Negative | Negative | 0.2 | Negative | Negative | - | Yellow | - |
| Negative | Negative | 0.2 | Negative | Negative | - | Yellow | - |
| Trace-lys | Negative | 0.2 | Negative | Negative | - | Yellow | - |
| Negative | Negative | 0.2 | Negative | Negative | - | Yellow | - |
| Negative | Negative | 0.2 | Negative | Negative | - | Yellow | - |
| Moderate | Negative | 0.2 | Negative | Negative | - | Yellow | - |
| Negative | Negative | 0.2 | Negative | Negative | - | Yellow | - |
| Negative | Negative | 0.2 | Negative | Negative | - | Yellow | - |
| Trace-lys | Negative | 0.2 | Negative | Large | - | Yellow | - |
| Negative | Negative | 0.2 | Negative | Negative | - | Yellow | - |
| Negative | Trace | 0.2 | Negative | Negative | - | Yellow | - |
| Large | Negative | 0.2 | Negative | Negative | - | Straw | - |
| Trace-lys | Negative | 0.2 | Negative | Negative | - | Yellow | - |
| Negative | Negative | 0.2 | Negative | Negative | - | Yellow | - |
| Negative | Negative | 0.2 | Negative | Negative | - | Yellow | - |
| Negative | Negative | 0.2 | Negative | Negative | - | Yellow | - |
| Negative | Negative | 0.2 | Negative | Negative | - | Yellow | - |
| Trace-lys | Negative | 0.2 | Negative | Negative | - | Yellow | - |
| Trace-int | Negative | 0.2 | Negative | Small | - | Yellow | - |
| Negative | Negative | 0.2 | Negative | Negative | - | Yellow | - |
| Negative | Negative | 0.2 | Negative | Negative | - | Yellow | - |
| Negative | Negative | 0.2 | Negative | Negative | - | Yellow | - |

| Sv (Milliliters) | Ictot () | S.G. (Sg) | U Ph() | 'Ssa ()' | 'Casts ()' | 'Epith ()' | 'Wbcu ()' |
|---|---|---|---|---|---|---|---|
| - | - | 1.006 | 8.5 | - | None Seen | Squam 0-1 | None Seen |
| - | - | 1.004 | 8.5 | - | None Seen | Squam 0-1 | None Seen |
| - | - | 1.01 | 8.5 | - | None Seen | Squam 0-1 | None Seen |
| - | - | 1.006 | 8.5 | - | None Seen | None Seen | None Seen |
| - | - | 1.006 | 8.5 | - | None Seen | None Seen | None Seen |
| - | - | 1.003 | 8.5 | - | None Seen | None Seen | None Seen |
| - | - | 1.003 | 8 | - | None Seen | Squam 0-1 | None Seen |
| - | - | 1.009 | 8.5 | - | None Seen | Squam 0-1 | None Seen |
| - | - | 1.01 | 8.5 | - | None Seen | Squam 0-1 | None Seen |
| - | - | 1.009 | 8.5 | - | None Seen | None Seen | None Seen |
| - | - | 1.003 | 7.5 | - | None Seen | Squam 2-5 | None Seen |
| - | - | 1.003 | 8.5 | - | None Seen | Squam 0-1 | 0-1 |
| - | - | 1.002 | 8.5 | - | None Seen | Squam 0-1 | None Seen |
| - | - | 1.006 | 8.5 | - | None Seen | None Seen | None Seen |
| - | - | 1.007 | 8.5 | - | None Seen | None Seen | None Seen |
| - | - | 1.002 | 8.5 | - | None Seen | Squam 0-1 | None Seen |
| - | - | 1.004 | 7.5 | - | None Seen | Squam 0-1 | None Seen |
| - | - | 1.005 | 8.5 | - | None Seen | Squam 0-1 | None Seen |
| - | - | 1.008 | 8.5 | - | None Seen | Squam 0-1 | 0-1 |
| - | - | 1.005 | 8.5 | - | None Seen | Squam 0-1 | 0-1 |
| - | - | 1.005 | 8.5 | - | None Seen | Squam 2-5 | None Seen |
| - | - | 1.005 | 8.5 | - | None Seen | Squam 0-1 | 5-Feb |
| - | - | 1.008 | 8.5 | - | None Seen | Squam 6-10 | None Seen |
| - | - | 1.015 | 8.5 | Negative | None Seen | None Seen | None Seen |
| - | - | 1.003 | 8.5 | - | None Seen | Squam 0-1 | None Seen |
| - | - | 1.01 | 8.5 | - | None Seen | None Seen | None Seen |
| - | - | 1.008 | 8.5 | - | None Seen | Squam 0-1 | None Seen |
| - | - | 1.004 | 8.5 | - | None Seen | Squam 0-1 | None Seen |
| - | - | 1.005 | 8.5 | - | None Seen | None Seen | None Seen |
| - | - | 1.007 | 8.5 | - | None Seen | Squam 0-1 | None Seen |
| - | - | 1.002 | 7.5 | - | None Seen | Squam 0-1 | None Seen |
| - | - | 1.005 | 8.5 | - | None Seen | Squam 0-1 | None Seen |
| - | - | 1.006 | 8.5 | - | None Seen | Squam 0-1 | None Seen |
| - | - | 1.007 | 8.5 | - | None Seen | None Seen | None Seen |
| - | - | 1.005 | 8.5 | - | None Seen | Squam 0-1 | None Seen |

| 'Rbcu ()' | 'Bctr ()' | 'Crys ()' |
|---|---|---|
| None Seen | 2+ | - |
| None Seen | Trace | None Seen |
| None Seen | 1+ | - |
| None Seen | None Seen | - |
| None Seen | Trace | - |
| None Seen | 2+ | None Seen |
| 0-1 | 4+ | None Seen |
| None Seen | 1+ | None Seen |
| None Seen | 1+ | None Seen |
| None Seen | Trace | - |
| None Seen | 2+ | None Seen |
| None Seen | None Seen | None Seen |
| None Seen | 1+ | None Seen |
| None Seen | Trace | None Seen |
| None Seen | None Seen | - |
| None Seen | 2+ | None Seen |
| None Seen | None Seen | None Seen |
| None Seen | Trace | None Seen |
| 5-Feb | Trace | None Seen |
| None Seen | Trace | None Seen |
| None Seen | 1+ | - |
| None Seen | 2+ | None Seen |
| None Seen | 1+ | None Seen |
| 0-1 | None Seen | - |
| 0-1 | 2+ | - |
| None Seen | 1+ | - |
| 0-1 | Trace | - |
| None Seen | 1+ | None Seen |
| None Seen | None Seen | - |
| None Seen | 1+ | - |
| None Seen | 3+ | None Seen |
| None Seen | 1+ | None Seen |
| None Seen | 1+ | None Seen |
| None Seen | 1+ | None Seen |
| None Seen | 1+ | None Seen |
